# Optimal iron concentrations for growth-associated polyhydroxyalkanoate biosynthesis in the marine photosynthetic purple bacterium *Rhodovulum sulfidophilum*

**DOI:** 10.1101/545053

**Authors:** Choon Pin Foong, Mieko Higuchi-Takeuchi, Keiji Numata

**Affiliations:** Biomacromolecules Research Team, RIKEN Center for Sustainable Resource Science, Wako, Saitama, Japan.

## Abstract

Polyhydroxyalkanoates (PHAs) are a group of natural biopolyesters that resemble petroleum-derived plastics in terms of physical properties but are less harmful biologically to the environment and humans. Most of the current PHA producers are heterotrophs, which require expensive feeding materials and thus contribute to the high price of PHAs. Marine photosynthetic bacteria are promising alternative microbial cell factories for cost-effective, carbon neutral and sustainable production of PHAs. In this study, *Rhodovulum sulfidophilum*, a marine photosynthetic purple nonsulfur bacterium with a high metabolic versatility, was evaluated for cell growth and PHA production under the influence of various media components found in previous studies. We evaluated iron, using ferric citrate, as another essential factor for cell growth and efficient PHA production and confirmed that PHA production in *R. sulfidophilum* was growth-associated under microaerobic and photoheterotrophic conditions. In fact, a subtle amount of iron (1 to 2 μM) was sufficient to promote rapid cell growth and biomass accumulation, as well as a high PHA yield during the logarithmic phase. However, an excess amount of iron did not enhance the growth rate or PHA productivity. Thus, we successfully confirmed that an optimum concentration of iron, an essential nutrient, promotes cell growth in *R. sulfidophilum* and also enhances PHA utilization.

## Introduction

Polyhydroxyalkanoates (PHAs) are a family of biopolyesters that are produced by a wide variety of microorganisms for the purpose of surviving in unfavorable growth and stress conditions; these molecules act as carbon and energy storage, redox regulators and cryoprotectants [1–3]. PHA is one of the well-known bioplastics (biobased and biodegradable) that has been extensively developed with the aims to overcome problems such as plastic solid wastes, harmful chemical substance leaching and dependence on nonrenewable fossil fuels. These are the main disadvantages of petroleum-derived synthetic plastics [4, 5].

However, the high price of PHAs has made them less competitive compared to conventional synthetic plastics due to the high cost of raw materials used in fermentation and downstream purification steps [6–8]. Heterotroph bacteria such as wild-type or engineered strains of *Cupriavidus necator* H16, *Alcaligenes latus*, *Pseudomonas putida* and *Escherichia coli* are the main workhorses for large-scale production of PHAs [9–11]. These heterotroph bacteria lack photosynthesis ability and thus require expensive carbon supplies to sustain their growth and PHA biosynthesis. On the other hand, photosynthetic microorganisms could produce their own foods by utilizing inexpensive and abundantly available resources such as sunlight, carbon dioxide and nitrogen and are thus potential next-generation microbial cell factories [12, 13]. Several studies have reported successful PHA production by photosynthetic bacteria such as cyanobacteria and purple bacteria [14, 15]. Additionally, PHA producers derived from marine bacteria and halophiles could also reduce production costs and lower contamination risk by using seawater as culture media [16–18]. A few research groups have been focusing on marine photosynthetic purple bacteria, which have the additional advantage of the ability to grow and biosynthesize PHA under microaerobic conditions [19, 20].

A marine photosynthetic purple nonsulfur bacterium, *Rhodovulum sulfidophilum*, is our target PHA producer, because it has high metabolic versatility [21, 22]. The presence of PHAs in this bacterium was first observed using Sudan Black B staining [23]. It has been previously determined that sodium chloride, vitamins, ammonium chloride, phosphate and carbon sources could affect PHA accumulation and cell growth of *R. sulfidophilum* [19, 20].

In this study, we evaluated another nutrient component, iron, which is required in many key biological reactions to support microbial growth and activity [24]. We examined the relationship between cell growth and PHA accumulation in a time-dependent manner by controlling the iron concentrations in the medium.

## Results

### Effect of iron on cell growth and PHA accumulation

Addition of 1 to 20 μM of iron in 520-Growth Medium (520-GM) enhanced the growth of *R. sulfidophilum* during the first three days of the culturing period compared to the 0 μM iron condition (Fig 1A-1C). In particular, low iron concentrations (1 to 5 μM) had positive effects on cell growth until Day-3, which corresponded to the logarithmic phase. The highest specific growth rate, 2.42 d^−1^ was achieved in the presence of 1 μM iron between Day-1 and Day-2 (Fig 1B). In the presence of iron, the specific growth rates decreased to negative values after Day-4, indicating that the cultures reached stationary phase faster than those in the 0 μM iron condition. An interesting observation was an extended logarithmic phase in the culture with 0 μM iron, even after the six day culture period. The same trend was observed for dry cell weights (DCWs) (Fig 1C). DCWs were relatively higher in lower iron concentration conditions until Day-3, while the DCW continued increasing even at Day-6 in cultures not treated with iron. The cultures reached maximum DCWs (~2.5 to 2.8 g/L) after entering stationary phase.

**Fig 1.**
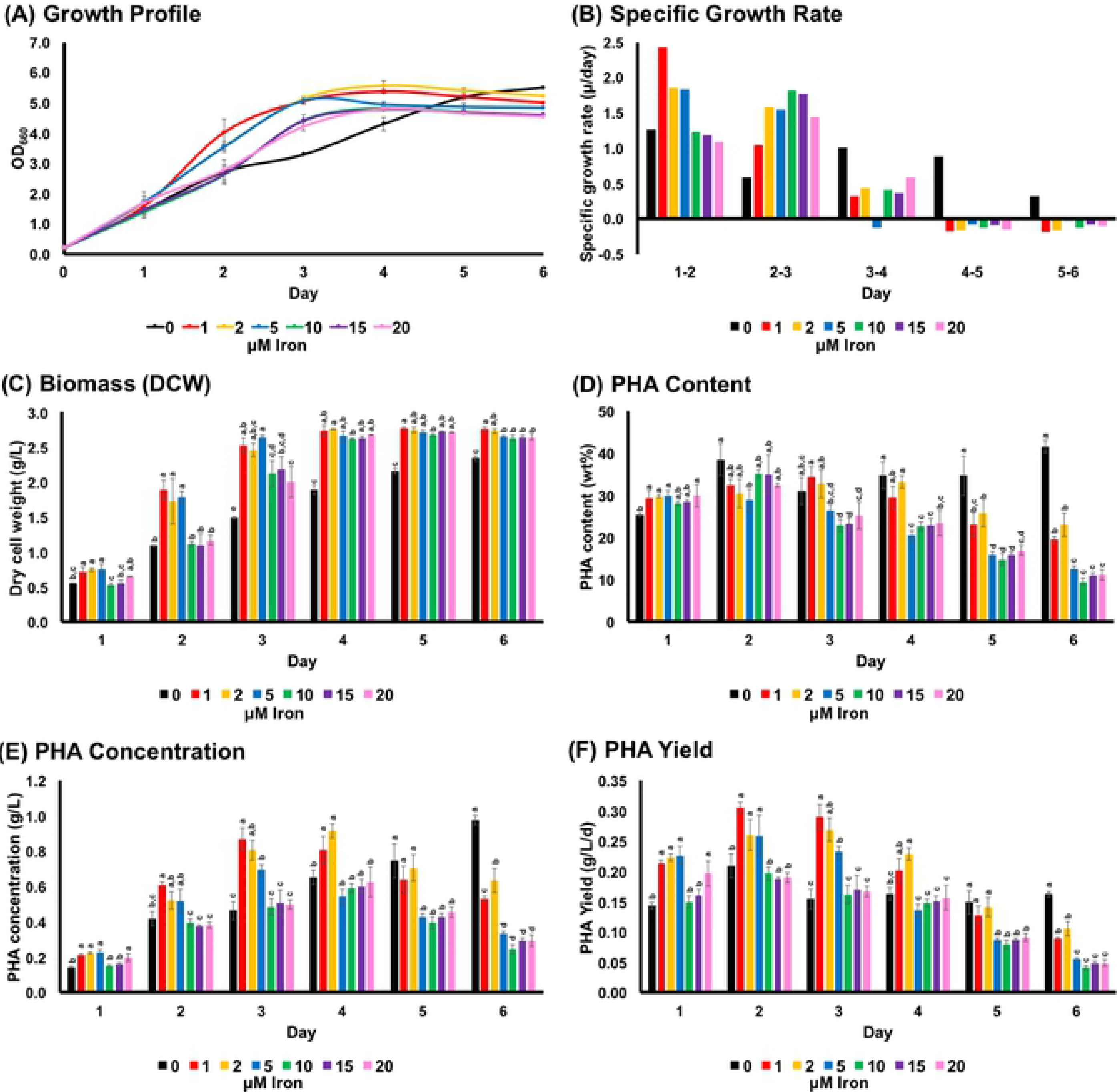
Effect of iron concentration (ferric citrate) on *R. sulfidophilum* under microaerobic and photoheterotrophic conditions. (A) cell growth (B) specific growth rate (C) biomass (D) PHA content (E) PHA concentration (F) PHA yield. Mean data accompanied by different superscripted letters are significantly different (Tukey’s HSD, p < 0.05).

It is also worth noting the discrepant results between optical density and DCW (Fig 1A and 1C) for some data points. For instance, there were no significant differences in term of DCW (2.6 to 2.8 g/L) between Day-4 and Day-5 (1 to 20 μM iron), but there were significant differences (data not shown) in OD_660_ (4.6 to 5.6). Indeed, this was clearly shown in the 0 μM iron culture at Day-6. This discrepancy could be due to the light scattering effect of PHA granules, which decreases the intensity of transmitted light in turbidity measurements [25], as has been shown in *Cupriavidus necator* H16 (wild-type) and its PHB-negative mutant (strain PHB^−^4).

On the other hand, the addition of iron elements, namely ferric citrate, did not improve the ability of *R. sulfidophilum* to accumulate PHAs in 520-GM (Fig 1D). Indeed, the PHA content gradually decreased in the presence of ≥5 μM iron after Day-2. Furthermore, the PHA content was significantly lower in cultures with ≥5 μM iron after Day-3. The only exception was the culture with 0 μM iron, in which the PHA content gradually increased during the six day culturing period. PHA concentrations were relatively higher in low iron concentration conditions (1 and 2 μM) until Day-4 because of the higher DCWs (Fig 1E). The low PHA contents for cultures with ≥5 μM iron, as shown in Fig 1D, lead to low PHA concentrations after Day-5. Only culture with 0 μM iron showed an increasing trend for PHA concentration because of increases in both DCW and PHA content during the culture period. Regarding PHA yield (i.e., PHA concentration per time of culture), *R. sulfidophilum* with 1μM iron achieved the highest yield of 0.29 g/L/d at Day-2 before the culture entered stationary phase (Fig 1F).

### Correlation between PHA accumulation and cell growth

As shown in Fig 1, iron concentrations affected both cell growth and PHA accumulation. However, there was no obvious relationship between iron concentration and the PHA accumulating ability of *R. sulfidophilum* in the first two days of culturing period. One of the possible explanations for the trend in PHA accumulation and utilization could be linked to the growth phase of *R. sulfidophilum*. PHA contents were high early in the cultivation period and then decreased gradually in the presence of iron (Fig 1D). Culture with 0 μM iron resulted in a continuous increase in DCW and PHA contents (Fig 1C and 1D). These results suggest that PHA was synthesized and accumulated during the logarithmic phase and then was actively utilized by cells during the late logarithmic and stationary phases. We could observe this relationship in the growth profile and in examining the PHA content versus the specific growth rate (Fig 2). PHA contents were relatively higher (>23 wt%) during culture periods with a positive SGR, but became relatively lower (<23 wt%) during periods with a negative SGR. Indeed, this was clearly shown in the 0 μM iron condition, where the PHA was not utilized by cells during the six day logarithmic phase.

**Fig 2.**
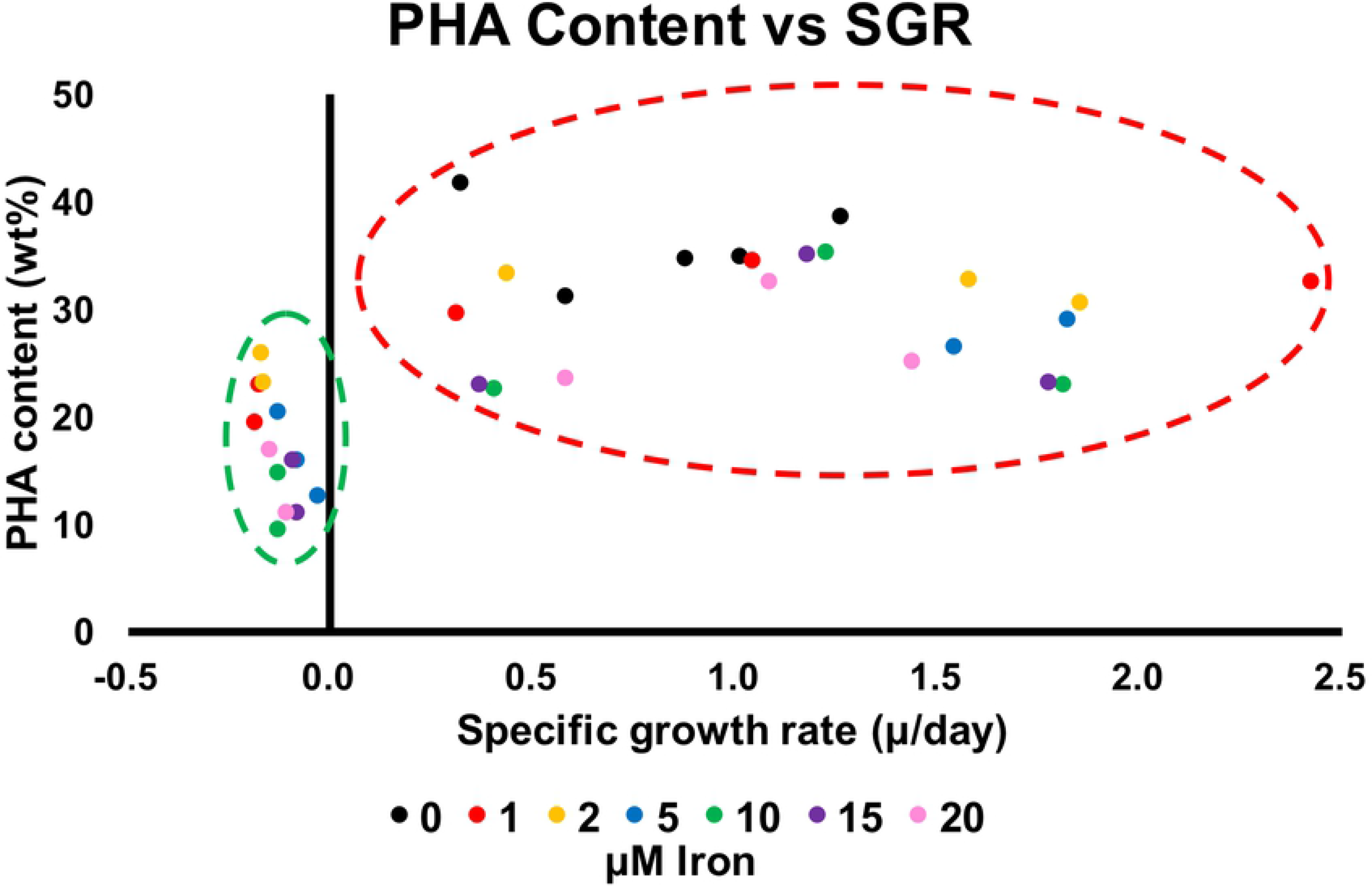
Correlation of PHA content with specific growth rate (SGR). PHA contents were relatively higher during periods of positive SGR (red-dashes circle), while PHA contents were relatively lower during periods of negative SGR (green-dashes circle).

## Discussion

Iron is one of the essential nutrients for major biological reactions and processes such as photosynthesis, nitrogen fixation, respiration, central metabolism and DNA repair [26]. The importance of iron in promoting cell growth of *R. sulfidophilum* was clearly shown in this study, where 1 μM iron was found to be optimal to achieve rapid cell growth and cell biomass accumulation in a shorter period of time compared to the 0 μM iron and excess iron conditions (Fig 1A-1C). On the other hand, high iron concentrations had a negative effect on cell growth in the logarithmic phase. Oversupply of iron could lead to generation of toxic free radicals or reactive oxygen species (ROS) in the cells via the Fenton/Haber-Weiss reaction in the presence of oxygen [27]. This could be the reason for the slower growth and cell biomass accumulation of *R. sulfidophilum* in the presence of ≥10 μM iron during the logarithmic phase.

Generally, accumulation of PHAs in most of the PHA-producing bacteria is triggered in conditions of unbalanced nutrients or in the absence of essential nutrients such as nitrogen, phosphorus, oxygen, sulfate, iron, magnesium, manganese, potassium or sodium [28] but with an excess of a carbon supply. Most of these bacteria synthesize PHA during the late logarithmic or stationary phases and are known as nongrowth-associated PHA producers [29]. However, there are always exceptions where PHA is synthesized during the logarithmic phase in growth-associated PHA producers, such as *Azohydromonas lata* (formerly known as *Alcaligenes latus*) [30]. Previous studies reported that *R. sulfidophilum* accumulates PHAs in a growth-associated manner during the logarithmic phase [20], and there is no enhancement of PHA production under nitrogen-limited conditions [19], which contrasts the findings for most of the nongrowth-associated PHA producers [29]. These observations were further supported by the results of this study. Indeed, growth-associated PHA producers are promising candidates for large-scale continuous PHA production (chemostat), which has advantages such as higher productivity, constant product quality and lower production costs [8].

Significant decreases in PHA content, PHA concentration and yield were detected after Day-2 or Day-3 of cultivation in the presence of 1 to 20 μM iron. Additionally, the PHA contents and PHA concentrations were also significantly lower in cultures with ≥5 μM iron after Day-3. This could be because of the competition between reduction of equivalents during both the PHA and hydrogen production processes and the utilization of PHA as a substrate for hydrogen production in *R. sulfidophilum* [31]. Hydrogen was produced in *R. sulfidophilum* during the late logarithmic and stationary phases [32]. Further investigations are needed to clarify this competitive relationship for reducing equivalents in *R. sulfidophilum*.

Chowdhury and coworkers reported that various medium compositions and concentrations could affect PHA accumulation and cell growth in *R. sulfidophilum*, such as sodium chloride, vitamins, ammonium chloride, phosphate and carbon sources. Both PHA production and cell growth were stimulated by addition of sodium chloride (3 wt%), ammonium chloride (5 mM) and phosphate. Removal of vitamins also greatly enhanced PHA accumulation but had no obvious effect on cell growth [20]. In this study, we successfully confirmed that iron is another essential nutrient that promotes cell growth in *R. sulfidophilum* and also enhances PHA utilization during the stationary phase. Thus, an optimum concentration of iron needs to be determined in order to achieve the highest PHA productivity. In summary, we conclude that 1 to 2 μM of iron in 520-GM was able to achieve the highest PHA yield of 0.26 to 0.29±0.02 g/L/d at Day-2 and Day-3 of cultivation (Fig 1F).

## Materials and Methods

### Media compositions and culture conditions

The marine phototrophic purple nonsulfur bacterium *Rhodovulum sulfidophilum* DSM1374/ATCC35886/W4 was obtained from the American Type Culture Collection (ATCC) biological resource center (BRC). For general cultivation purpose, *R. sulfidophilum* was maintained on marine agar or marine broth (BD Difco, New Jersey, USA) under microaerobic condition at 30°C with continuous far-red LED light (730 nm, 20 to 30 Wm^−2^).

A different rich medium, 520-Growth Medium (520-GM), was used for PHA accumulation purposse. The 520-GM contained the following components per liter: 0.5 g KH_2_PO_4_, 0.25 g CaCl_2_⋅2H_2_O, 3.0 g MgSO_4_⋅7H_2_O, 0.68 g NH_4_Cl, 20 g NaCl, 3.0 g sodium L-malate, 3.0 g sodium pyruvate, 0.4 g yeast extract, 2 mg vitamin B12 and micronutrients including 70 μg ZnCl_2_, 100 μg MnCl_2_⋅4H_2_O, 60 μg H_3_BO_3_, 200 μg CoCl_2_⋅6H_2_O, 20 μg CuCl_2_⋅2H_2_O, 20 μg NiCl_2_⋅6H_2_O and 40 μg Na_2_MoO_4_⋅2H_2_O. The pH of the medium was adjusted to 7.0 before autoclave sterilization.

Different concentrations of ferric citrate, ranging from 0 to 5.0 mg/L (1 to 20 μM of iron) were added into 520-GM to evaluate the effects on cell growth and PHA accumulation. A one-stage cultivation strategy was employed for PHA production, where the *R. sulfidophilum* was first precultured in 520-GM until the OD_660_ value reached approximately 2.0 (logarithmic phase) before being transferred to 520-GM with a final OD_660_ value of 0.2 after dilution. A total of 15 mL 520-GM in a 15 mL conical centrifuge tube was used, and the cultures were incubated for six days under microaerobic conditions at 30°C with continuous far-red LED light (730 nm, 20 to 30 Wm^−2^). Cells were harvested by centrifugation at 9,000 × *g* for 10 min, washed with distilled water once, kept at −80°C overnight and then freeze-dried for 24 h.

### Growth profile and cell biomass analyses

The growth profile of *R. sulfidophilum* was measured using a UV/Vis-spectrophotometer at an absorbance of 660 nm with a 24-h interval. Specific growth rates (μ/day) were calculated based on the slope of the growth curve [33]. Total cell biomass (g/L) was determined based on dry cell weight of the samples.

### PHA quantification

Approximately 2 mg of lyophilized cells were subjected to methanolysis [1 mL chloroform and 1 mL methanolysis solution (methanol:sulfuric acid, 85:15 v/v)] at 100°C for 140 min. After cooling to room temperature, 1 mL of phosphate buffer (pH 8) was added to the reaction mixture, which was then vortex-mixed and neutralized with 0.5 N NaOH. The lower chloroform layer was transferred to a glass tube containing sodium sulfate anhydrous to remove water content. The PHA content and composition were determined using gas chromatography-mass spectrometry (GC-MS) machine (GCMS-QP2010 Ultra, Shimadzu, Kyoto, Japan) equipped with a 30 m × 0.25 mm DB-1 capillary gas chromatography column (Agilent Technologies, CA, USA). For analysis, a 1 μL volume of sample was injected with helium as a carrier gas (3.30 mL min^−1^). The following temperature program was used to separate ethyl esters: 45°C for 1 min, temperature ramp of 7°C per min to 117°C. The interface and ion source temperatures were 250°C and 230°C, respectively.

### Statistical analysis

The Statistical Package for the Social Sciences (SPSS) software version 22 (IBM Corp. Released 22.0.0.0, New York, USA) was used for all analyses. Statistically significant differences between groups were determined by one-way analysis of variance (ANOVA) assessments with Tukey post hoc tests, where a p-value <0.05 indicated a significant difference.

## Acknowledgements

This work was supported by the ImPACT Program of the Council for Science, Technology and Innovation (Cabinet Office, Government of Japan) and JST ERATO (Grant Number JPMJER1602), Japan.

